# The Neural Correlates of Semantic Control Revisited

**DOI:** 10.1101/2020.07.15.204990

**Authors:** Rebecca L. Jackson

**Affiliations:** MRC Cognition & Brain Sciences Unit, University of Cambridge, 15 Chaucer Road, Cambridge, CB2 7EF

**Keywords:** semantic cognition, control, ALE meta-analysis, executive processing, semantic control

## Abstract

Semantic control, the ability to selectively access and manipulate meaningful information on the basis of context demands, is a critical component of semantic cognition. The precise neural correlates of semantic control are disputed, with particular debate surrounding parietal involvement, the spatial extent of the posterior temporal contribution and network lateralisation. Here semantic control is revisited, utilising improved analysis techniques and a decade of additional data to refine our understanding of the network. A meta-analysis of 876 peaks over 121 contrasts illuminated a left-focused network consisting of inferior frontal gyrus, posterior middle temporal gyrus, posterior inferior temporal gyrus and dorsomedial prefrontal cortex. This extended the temporal region implicated, and found no parietal involvement. Although left-lateralised overall, relative lateralisation varied across the implicated regions. Supporting analyses confirmed the multimodal nature of the semantic control network and situated it within the wider set of regions implicated in semantic cognition.

**Highlights:**

➢ A multimodal semantic control network was delineated with formal meta-analyses
➢ Semantic control recruits inferior and medial frontal and posterior temporal cortex
➢ A large extent of posterior temporal cortex was implicated and no parietal regions
➢ Semantic control is left-lateralised but regions show differential lateralisation
➢ The semantic control regions were situated in the context of the wider semantic network

## Introduction

Semantic cognition is comprised of two distinct, yet interacting elements; semantic representation and semantic control, a distinction that forms the basis of the Controlled Semantic Cognition framework (Jefferies, 2013; Lambon Ralph et al., 2017). Semantic representation is the extraction and storage of the underlying structure within the environment, abstracting conceptual knowledge across learning episodes, sensory modalities and task contexts (Lambon Ralph et al., 2017). This representation element is impaired in semantic dementia (SD); the gradual loss and blurring of such representations resulting in a loss of the ability to comprehend words, pictures and objects of all categories and across all sensory input modalities (Hodges and Patterson, 2007; Patterson et al., 2007). Semantic representation critically depends on the interaction between the modality-specific spoke regions distributed throughout the cortex and the multimodal hub region in the ventral anterior temporal lobe (ATL; Pobric et al., 2007; Binney et al., 2010; Pobric et al., 2010; Acosta-Cabronero et al., 2011; Abel et al., 2015; Lambon Ralph et al., 2017). By mediating between distributed sensorimotor input and output representations, the ATL can extract and represent the underlying multimodal semantic structure across learning episodes (Rogers et al., 2004; Rogers and McClelland, 2004; Patterson et al., 2007; Lambon Ralph et al., 2017; Jackson et al., submitted).

Less research pertains to the second element of semantic cognition; semantic control. Semantic control is the ability to selectively access and manipulate meaningful information based on context demands and is also hypothesised to be multimodal (Jefferies, 2013). In semantic aphasia (SA), cerebrovascular accident to frontal or temporoparietal cortex affects this ability independently of the stored representations (Jefferies and Lambon Ralph, 2006). Intriguingly, frontal and posterior lesions present with the same behavioural profile, suggesting a distributed network underlying semantic control, including inferior frontal and posterior temporal and/or inferior parietal regions (Jefferies, 2013). At odds with the dual foci of these lesion patterns, early imaging results focused on the inferior frontal gyrus (IFG) alone, particularly pars triangularis (e.g., Thompson-Schill et al., 1997; Wagner et al., 2001). However, a meta-analysis contrasting more over less controlled semantics identified additional posterior involvement in accord with the neuropsychological data. Specifically, Noonan et al., (2013) identified areas with high activation likelihood in a left-focused network, including posterior middle temporal gyrus (pMTG), inferior parietal cortex, anterior cingulate and anterior MTG, as well as bilateral IFG and dorsomedial prefrontal cortex (dmPFC). This added greater spatial precision to the regions theorised to causally underpin semantic cognition based on the neuropsychological results.

Since this meta-analysis of semantic control (Noonan et al., 2013), control processes have gained increasing recognition and many more studies have directly manipulated the level of control required within semantic tasks. Simultaneously, imaging protocols have improved in a multitude of ways, increasing spatial specificity and statistical power, as well as gaining better coverage across cortical regions critical for semantic cognition (e.g., Feinberg and Setsompop, 2013; Ugurbil et al., 2013; Halai et al., 2014). This additional high quality data, combined with improvements in meta-analytical tools, (which now support more appropriate FWE thresholding procedures; Eickhoff et al., 2012; Eickhoff et al., 2017), gives an opportunity to return to the meta-analytic approach to provide an updated map, refining our understanding of the underlying neural correlates. Critically, this revision could help resolve a set of remaining puzzles as to the precise cortical anatomy of semantic control.

Debate as to the neural correlates of semantic control surrounds three open issues: 1) the involvement of inferior parietal cortex, 2) the spatial extent of lateral posterior temporal cortex, and 3) the lateralisation of the semantic control network. The role of inferior parietal regions is disputed, both in semantic cognition generally and semantic control specifically (e.g., Binder and Desai, 2011; Humphreys and Lambon Ralph, 2014). Noonan et al., (2013) identified one cluster implicating a region at the border of dorsal angular gyrus (AG) and inferior parietal sulcus (IPS), postulated to be a domain-general executive control region. Additionally, a smaller cluster in ventral AG showed greater involvement in harder semantic cognition, which was considered puzzling due to the overlap with the default mode network (expected to show greater activation, or relatively less deactivation, for easier tasks and rest; Buckner et al., 2008). This functional division between IPS and AG was supported by a large cross-domain meta-analysis showing ventral AG deactivation for ‘automatic semantics’ (Humphreys and Lambon Ralph, 2014). Furthermore, multiple functional regions may exist within the AG and task involvement may not map neatly onto the anatomical divisions (Caspers et al., 2008; Seghier, 2013). In combination with the lack of spatial precision of SA patients’ ‘temporoparietal’ damage, these findings have led to persisting uncertainty as to the location of a possible inferior parietal semantic control region, with authors labelling this region using vague terms, such as ‘IPL/IPS’ (e.g., Jefferies, 2013; Jackson et al., 2016), or focusing on IPS alone (e.g., Davey et al., 2016; Lambon Ralph et al., 2017; Hoffman, 2018). Can a meta-analysis with additional data provide evidence adjudicating the role of these inferior parietal regions in semantic control?

The second critical debate regards the spatial extent of posterior lateral temporal cortex involvement. Noonan et al., (2013) specifically highlighted the involvement of the pMTG in semantic control. However, whilst posterior lateral temporal activity is often found when assessing semantic control, the precise region implicated can wander into contiguous gyri (Thompson-Schill et al., 1997; Snyder et al., 2011; Rodd et al., 2012). Is the focus on pMTG in the literature an accurate depiction of the posterior temporal regions responsible for semantic control? The third unresolved issue is the laterality of semantic control. Noonan (2013) identified greater involvement of the left hemisphere with some activation of right frontal cortex. Although rarely studied, right hemisphere damage appears to produce a qualitatively similar, yet quantitatively reduced, control impairment (Thompson et al., 2016). Would greater power result in a more bilateral profile with involvement of right temporal and parietal cortex?

An updated meta-analysis will determine the regions consistently implicated in semantic control, helping address these puzzles: which (if any) parietal regions are implicated, what is the spatial extent of posterior temporal involvement, and is the network strongly left-lateralised throughout? Furthermore, the additional data makes it possible to independently assess the regions implicated in semantic control with visual and auditory stimuli and directly contrast them, testing whether the network is multimodal as hypothesised within the Controlled Semantic Cognition framework. Whilst Noonan et al., (2013), argued for the clear need to perform this test, it was not possible with only 9% of the studies employing auditory stimuli. Additionally, through assessment of the regions implicated in semantic cognition more generally, we can assess how this semantic control network is situated within the wider context of semantic cognition areas.

## Materials & Methods

Meta-analyses were employed to ask 1) which regions are involved in semantic control, 2) are the same regions involved in semantic control with visual and auditory stimuli, and 3) how do semantic control areas relate to those implicated in general semantic cognition.

### Inclusion and Exclusion Criteria for Semantic Control

The inclusion criteria were based on those instantiated in Noonan et al., (2013), focusing on PET and fMRI studies manipulating the amount of semantic control required by contrasting more controlled (and harder) semantic cognition over less controlled (and easier) semantic cognition. However, some additional restrictions were possible with the increased number of studies assessed (or necessary due to additions to the literature in recent years). All studies were required to report peak differences in univariate activation values in a standard space (Talairach/MNI) in a peer-reviewed English language article. Tasks meeting the inclusion criteria comprised manipulations of homonym ambiguity, competitor interference, association strength, semantic violations, meaning dominance and alternative uses of an object. Where present, multiple distinct contrasts were included from a single study. Studies were excluded if focused on patients, gender differences, priming or cueing, bilingualism, developmental semantics, episodic memory, sleep consolidation, learning novel semantics or ageing. Participants were required to be between 18 and 65. Contrasts of different stimuli types (e.g., animals vs. tools, metaphoric vs. literal sentences), manipulations of psycholinguistic variables (e.g., imageability), manipulations of attention or multimodal integration, changes in perception or timing and manipulations of sentences order or syntactic violations were not considered to fit these criteria. Manipulations of executive control demands (e.g., go *vs*. no go) with meaningful stimuli were excluded as the core contrast is not focused on semantic demands. Comparisons of participants with differing ability or correct *vs*. incorrect trials were also excluded.

### Inclusion and Exclusion Criteria for General Semantics

The same inclusion and exclusion criteria were used for the general semantic contrast except those relating to the nature of the contrast. Studies were only included in this contrast if they compared more > less semantic cognition, either by contrasting a semantic with a non- (or less) semantic task or meaningful (or known) with meaningless (or unknown) stimuli (including intelligibility assessments). This did not include comparison of high and low familiarity (as either could elicit more semantic processing) or imageability (as both concrete and abstract items require semantic processing and the nature of this processing may differ in numerous ways). Studies recruiting rest (or fixation) as a baseline were excluded due to the known issues in contrasting semantics to low-level baselines, whereby key regions may be missed due to the high level of semantic processing present during rest (Visser et al., 2010). In addition to a substantial update to the timeframe of study inclusion, the present approach differs from the prior meta-analysis by Binder et al., (2009) on two critical aspects: 1) both verbal and nonverbal stimuli are included as semantic cognition is considered inherently multimodal, and 2) it is not required that the baseline control task be at least as difficult as the semantic task (as this induces a difficulty difference) but merely that a high level baseline be employed.

### Identifying Studies

The studies assessed for inclusion were sourced from prior meta-analyses of semantic control and semantic cognition; Noonan et al., (2013), Humphreys & Lambon Ralph (2014), Binder et al., (2009), and Rice et al., (2015a) and a Web of Science (formerly Web of Knowledge; https://clarivate.com/products/web-of-science/) search designed to extend the timeframe of inclusion. Whilst Noonan et al., (2013) included a limited number of studies in 2009, Binder al.’s (2009) coverage ended in 2007. Therefore, to ensure identification of all studies relevant to either contrast, the search was conducted from the start of 2008 until the time of assessment (19^th^ June 2019). This search employed the same search terms as Noonan et al., (2013); ‘semantic’ or ‘comprehension’ or ‘conceptual knowledge’ in conjunction with imaging terms ‘fMRI’ or ‘PET’. Due to the large number of studies identified in this search, a set of exclusion terms related to the exclusion criteria were included; patient, priming, disorder, dementia, aging, ageing, bilingual, meta-analysis, multivariate. Overall, 2052 studies were assessed for the fit to the inclusion criteria; 1835 from Web of Knowledge and 217 from prior meta-analyses. This resulted in 121 contrasts including 876 peaks for semantic control and 423 contrasts including 3860 peaks for general semantic cognition. The semantic control meta-analysis was split into visual and auditory verbal semantic control on the basis of the modality of the stimuli. The small number of contrasts with nonverbal stimuli were excluded as these were only present in the verbal condition and stimulus type may have a different effect to stimulus modality. Auditory verbal semantic control included 175 peaks across 23 contrasts and visual verbal semantic control included 649 peaks in 87 contrasts. All data included are provided in Supplementary Tables 1 and 2.

### Meta-Analysis Method

The meta-analyses were Activation Likelihood Estimates performed in GingerAle version 3.02 (available at http://www.brainmap.org/software.html#GingerALE; Eickhoff et al., 2009; Eickhoff et al., 2012; Turkeltaub et al., 2012; Eickhoff et al., 2017). All peaks were converted to MNI standard space within GingerAle and analyses performed in MNI space. Each contrast is used to construct a Model Activation map, which includes a Gaussian curve centred on each peak (Eickhoff et al., 2009; Turkeltaub et al., 2012). The full width at half maximum (FWHM) of the Gaussian is determined based on the sample size of the study, resulting in smoothing reflecting the uncertainty of the peak location (Eickhoff et al., 2009). A larger, tighter curve is employed around peaks with a larger sample size. No additional smoothing was performed. The union of the Model Activation maps from each contrast is the Activation Likelihood Estimation (ALE) map which reflects the agreement in identification of peaks across studies (Eickhoff et al., 2009; Turkeltaub et al., 2012). A p-value image is constructed based on the values of each voxel across the set of Model Activation maps reflecting the likelihood of finding that voxel in a study and then thresholded. Cluster-level permutation testing was used to control for the family-wise error rate as recommended by Eickhoff et al., (2012; 2017). Permutation testing is used to determine the size of cluster which would appear under the null hypothesis in only 5% of datasets. Removing clusters that fail to meet this size criterion applies FWE-correction at the cluster level. The null distribution may be generated within GingerAle using Monte-Carlo simulation where foci are randomly placed throughout the grey matter template and the largest cluster size recorded. All contrasts were performed with voxel-level thresholding at a p-value of .001 and cluster-level FWE-correction with a p-value of .001 over permutation testing with 10000 permutations.

These methodological details provide additional improvements upon Noonan et al., (2013) as the FWE-cluster correction is considered a more rigorous thresholding method and the individual-subject based smoothing method allowing the certainty based on sample size to be taken into account instead of simply applying a large, consistent amount of smoothing, has been demonstrated to improve meta-analyses (Eickhoff et al., 2009). Indeed, the FDR-based permutation testing performed in Noonan et al., (2013) was implemented with a known error in GingerAle further affecting the correction for multiple comparisons (Eickhoff et al., 2017).

Individual meta-analyses were used to construct activation likelihood maps for semantic control, visual verbal semantic control, auditory verbal semantic control and general semantics. The resulting maps for visual and auditory semantic control were directly contrasted within GingerAle, which allows identification of regions significantly more likely to be activated in each condition and a conjunction result; areas activated in both conditions (expressed as an ALE map; Eickhoff et al., 2011). These contrast analyses involve a subtraction of the thresholded maps and construction of a thresholded Z-score map for ease of interpretation. Contrast analyses were assessed with a p-value of .001, 10000 permutations and a minimum cluster volume of 20mm^3^. The results of all analyses are available online as mask files (https://github.com/JacksonBecky/SemanticControlMetaA).

## Results

### Semantic Control Regions

The areas identified in the semantic control contrast are displayed in Figure 1. The peak coordinates are listed in Table 1. The largest and strongest cluster encompasses the entire left IFG (including pars triangularis, pars orbitalis and pars opercularis) with some involvement of the insula and left orbitofrontal cortex. The strongest activation likelihood is within pars triangularis. A second cluster is focused on left posterior lateral temporal cortex, with activation covering a large portion of pMTG and posterior inferior temporal gyrus (pITG), as well as the edge of the fusiform gyrus. Activation likelihood peaks are found within both pMTG and pITG. A bilateral dmPFC cluster with a left-sided focus includes supplementary and pre-supplementary motor areas. Two clusters are identified within the right IFG; one centred on pars triangularis and a more ventral cluster including both pars orbitalis and the insula. Consistent with both the prior meta-analysis and the neuropsychological data, the present results indicate involvement of a distributed, left-dominant network of inferior frontal and lateral posterior temporal cortices in semantic control. Building upon this, the current results highlight the contribution of posterior temporal cortex outside the MTG, within the ITG. Unlike the previous semantic control meta-analysis, no parietal, ventromedial prefrontal or anterior temporal regions were found to be involved in semantic control.

**Figure 1.**
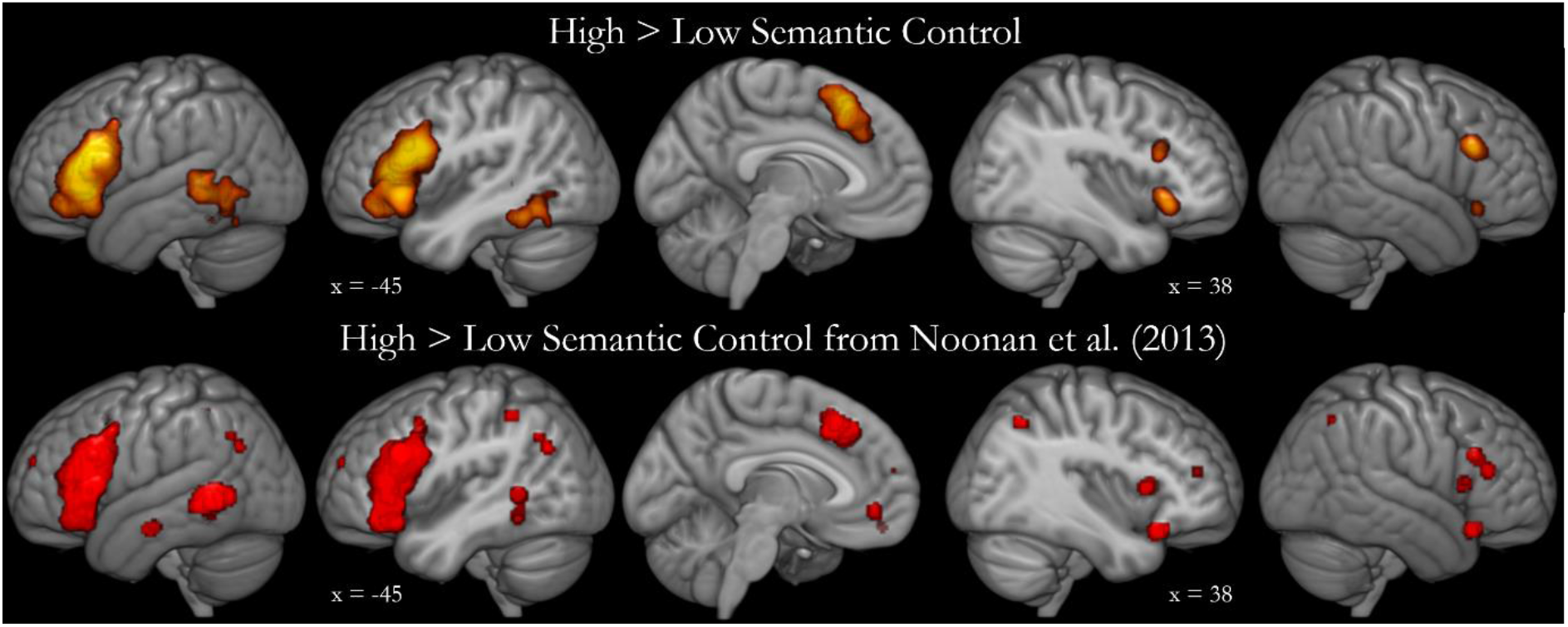
Results of the meta-analysis contrasting high > low semantic control. Top: Activation likelihood estimate map from the new extended analysis of semantic control based on 876 peaks from 121 contrasts comparing high > low semantic control. Activation likelihood is significant at a voxel-level of .001 and an FWE-corrected cluster-level of .001. Bottom: Regions identified as responsive to high > low semantic control in Noonan et al., 2013.

**Table 1.**
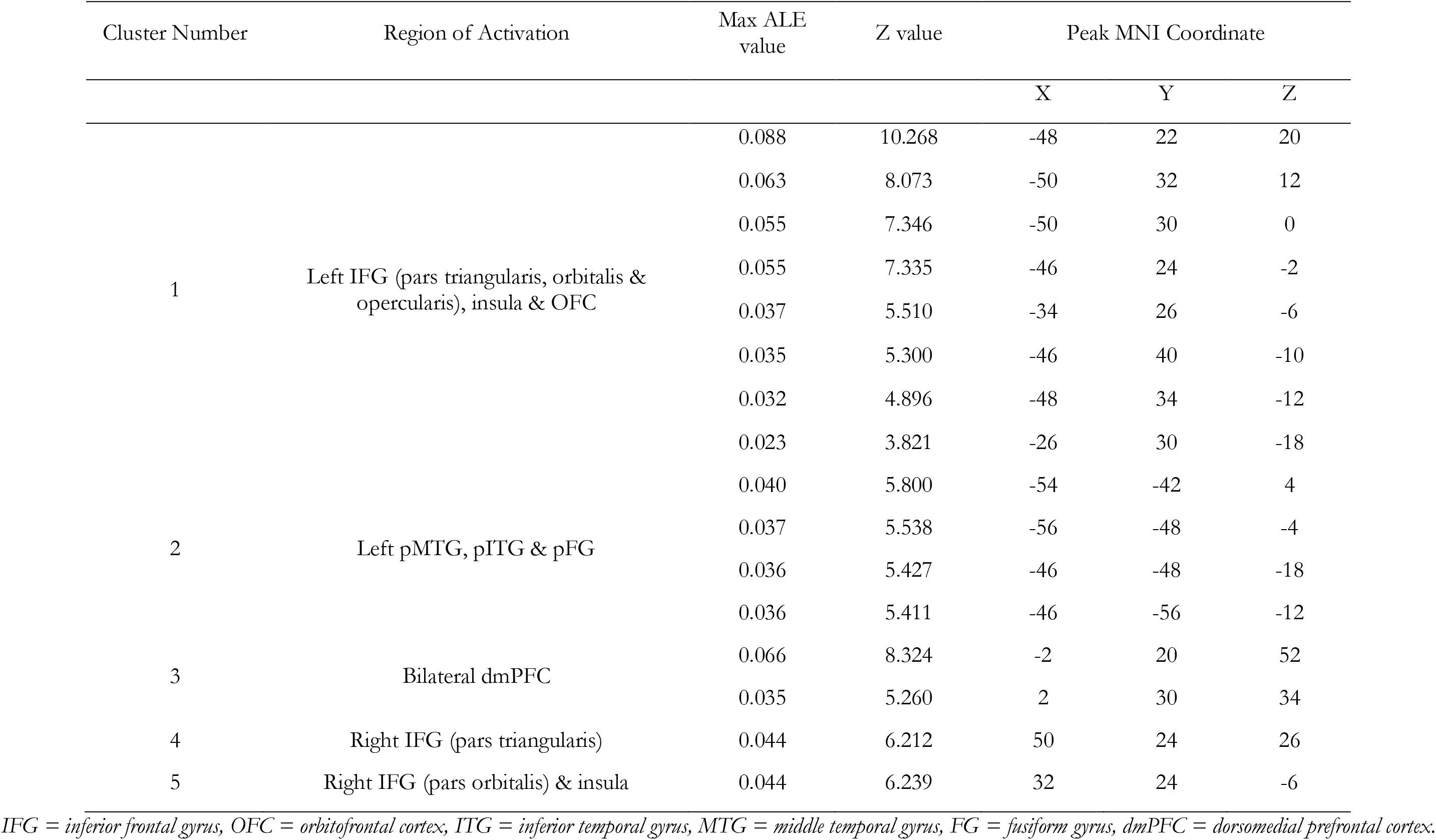
Semantic control activation likelihood.

### Visual and Auditory Semantic Control

The regions involved in semantic control with auditory and visual verbal stimuli are displayed in Figure 2, with peaks of activation likelihood listed in Table 2. Visual semantic control includes all of the clusters identified within the full semantic control analysis (left IFG and insula, left pMTG and pITG, bilateral dmPFC and two right IFG clusters). Although fewer contrasts were included, the auditory semantic control contrast highlights the two largest regions of involvement; left IFG (pars triangularis and opercularis) and posterior lateral temporal cortex, here focused on pITG. A conjunction analysis demonstrated overlap between the auditory and visual semantic control maps within left IFG and posterior temporal cortex (specifically in the pITG). Contrasting auditory and visual semantic control failed to identify any regions with greater involvement in either visual or auditory studies. Thus, the distributed network of inferior prefrontal and posterior temporal regions is implicated in semantic control regardless of input modality.

**Figure 2.**
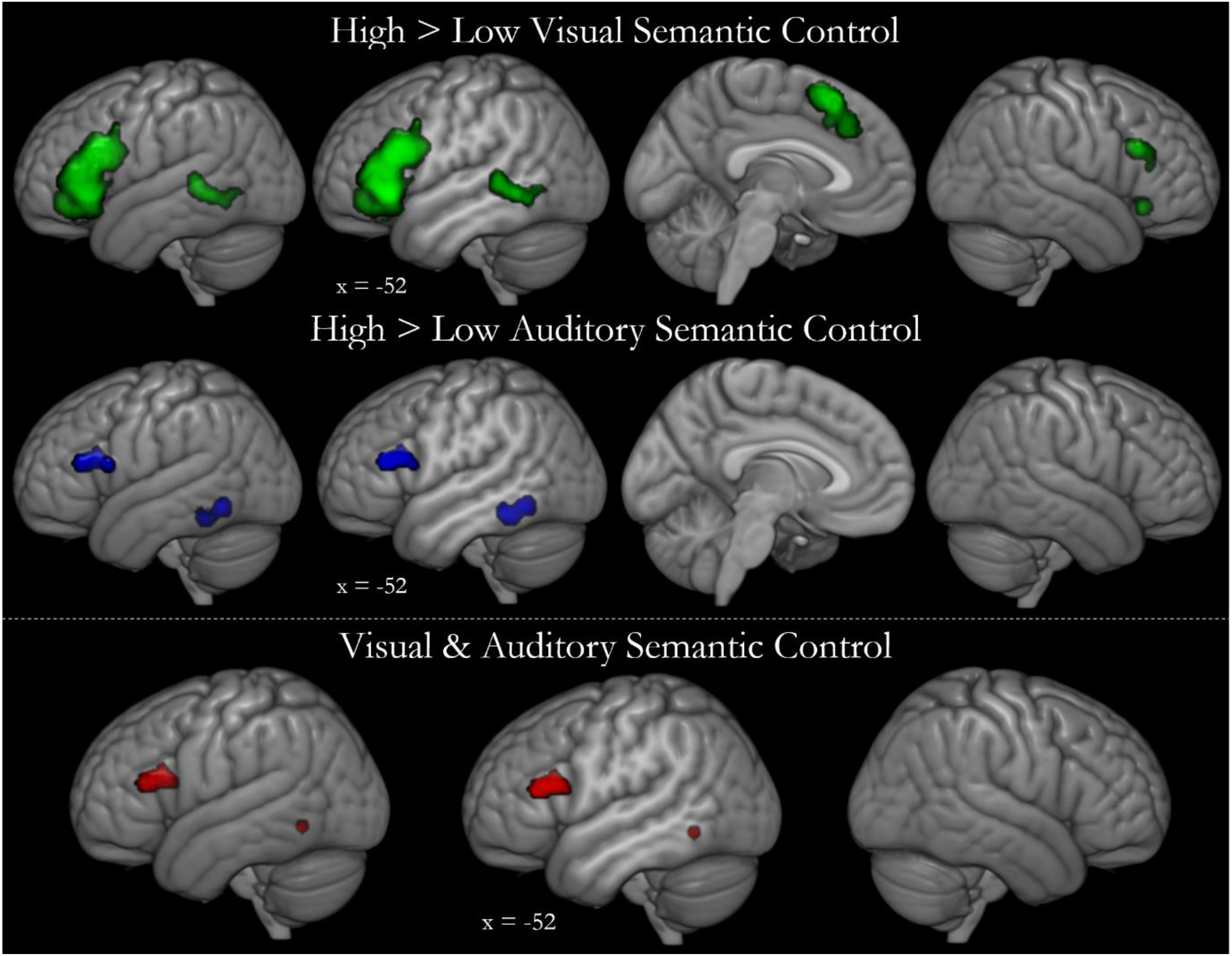
The multimodal semantic control network. Top: The activation likelihood estimate map for visual semantic control, based on 649 peaks from 87 contrasts, shown in green. The activation likelihood estimate map for auditory semantic control, based on 175 peaks from 23 contrasts, shown in blue. Activation likelihood is significant at a voxel-level of .001 and an FWE-corrected cluster-level of .001. Bottom: Contrasting visual and auditory semantic control allows visualisation of the conjunction of the two thresholded maps (the activation likelihood of the intersection is shown in red). Direct contrasts of auditory and visual semantic control did not result in any significant clusters.

**Table 2.**
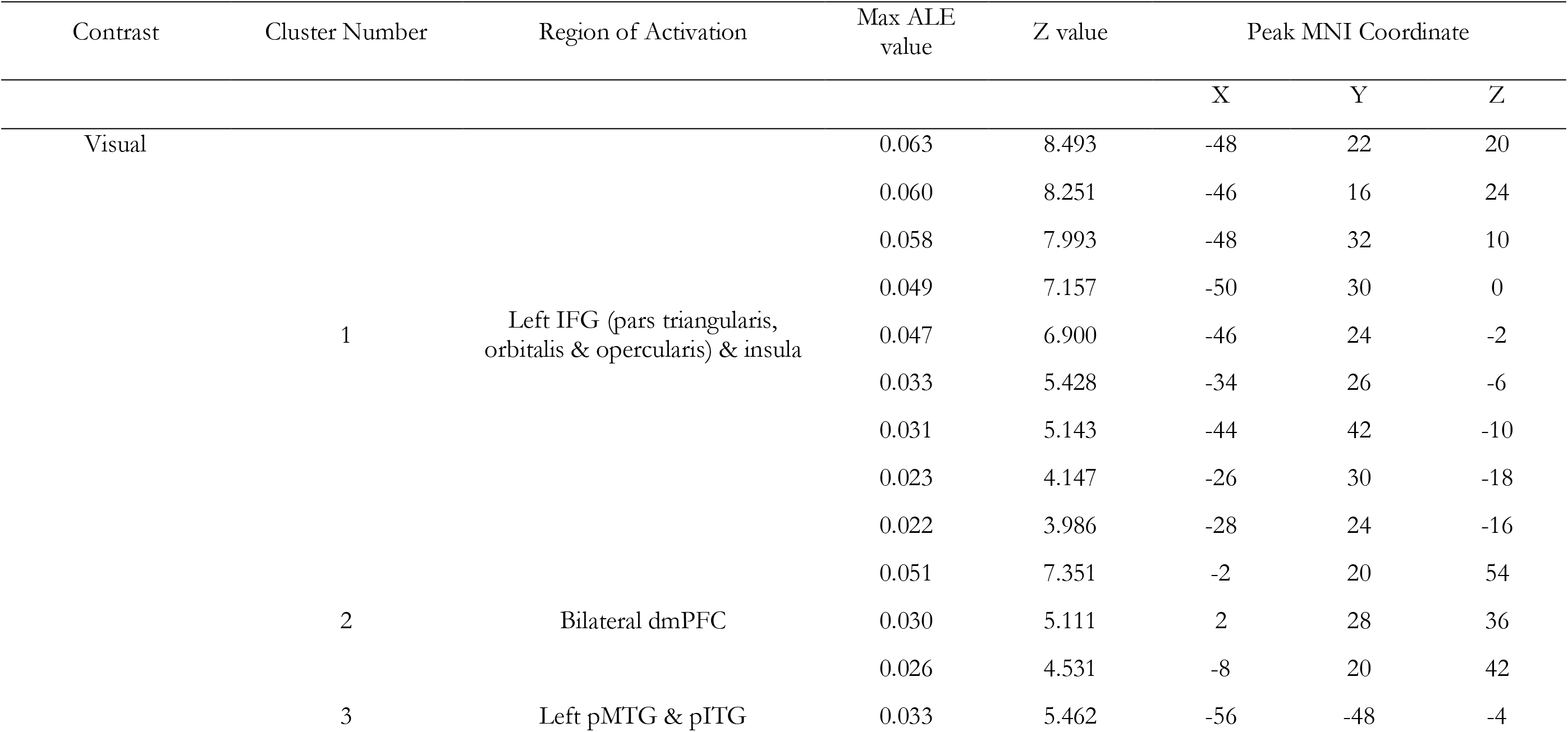
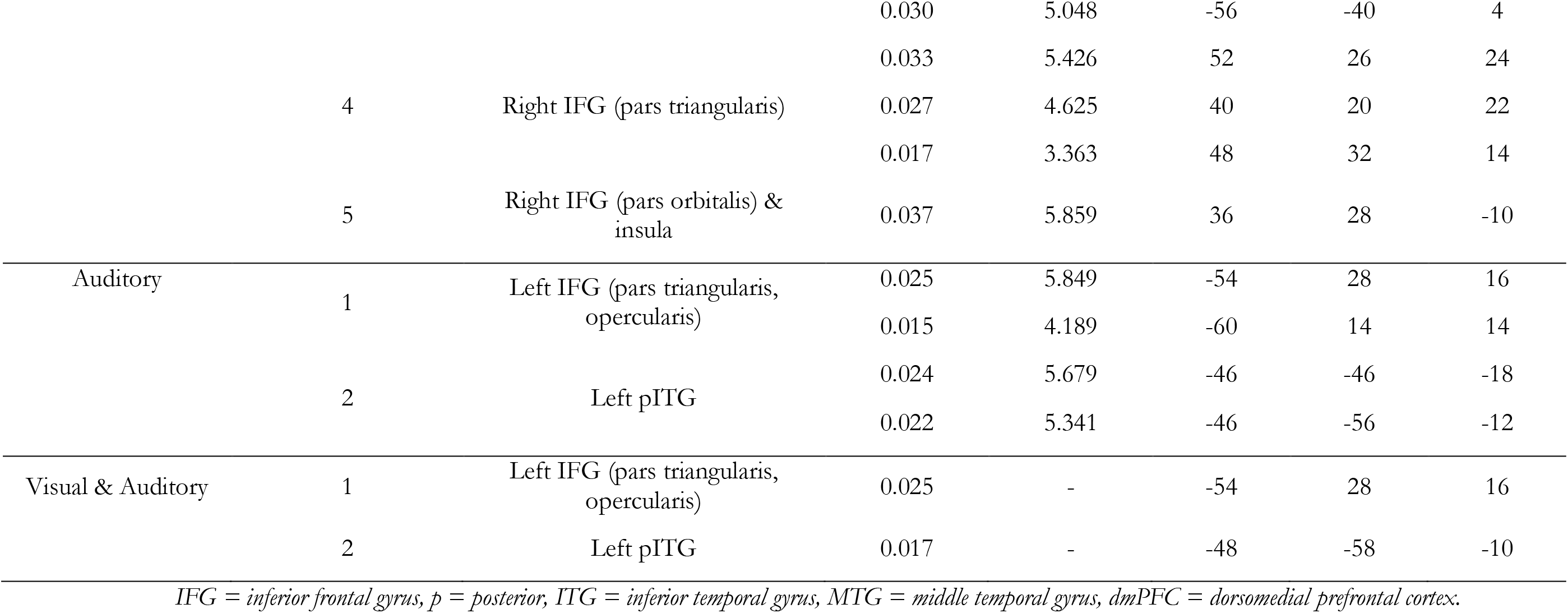
Auditory and visual semantic control activation likelihood.

### Semantic Control in the Wider Semantic Network

Contrasting semantic tasks and meaningful stimuli with baseline tasks and meaningless stimuli allowed identification of the larger network of regions implicated in semantic cognition (see Figure 3 & Table 3). One large cluster traversed left frontal, temporal and parietal cortex, covering the length of the MTG. This cluster subsumed IFG and included ventral ATL, pITG, superior temporal gyrus, hippocampus, insula and the inferior parietal cortex, including the AG. Additional clusters focused on bilateral dmPFC, right superior and middle temporal gyri, right IFG and insula and a left-focused posterior cingulate region. This pattern is in high accordance with the known architecture of the semantic system and the results of prior meta-analyses of semantics (Binder et al., 2009; Humphreys and Lambon Ralph, 2014; Rice et al., 2015a) and overlaps the regions implicated in semantic control. Specifically, all regions implicated in semantic control are found in the meta-analysis of general semantic cognition, except the right dorsal IFG cluster. This may require extremely controlled processing or may show a domain general executive pattern and therefore be lost in the comparison with other domains. All of the left frontal semantic regions are implicated in semantic control specifically, yet the temporal lobe shows a more complex pattern. Whilst a large portion of the left temporal lobe is implicated in semantic cognition, the majority is responsible for semantic representation with only the most posterior inferior and middle temporal regions implicated in control. This control area is flanked by temporal and parietal areas responsible for representation, which may provide some clues as to the interaction between, and organisation of, control and representation processes within the wider network (see Discussion).

**Figure 3.**
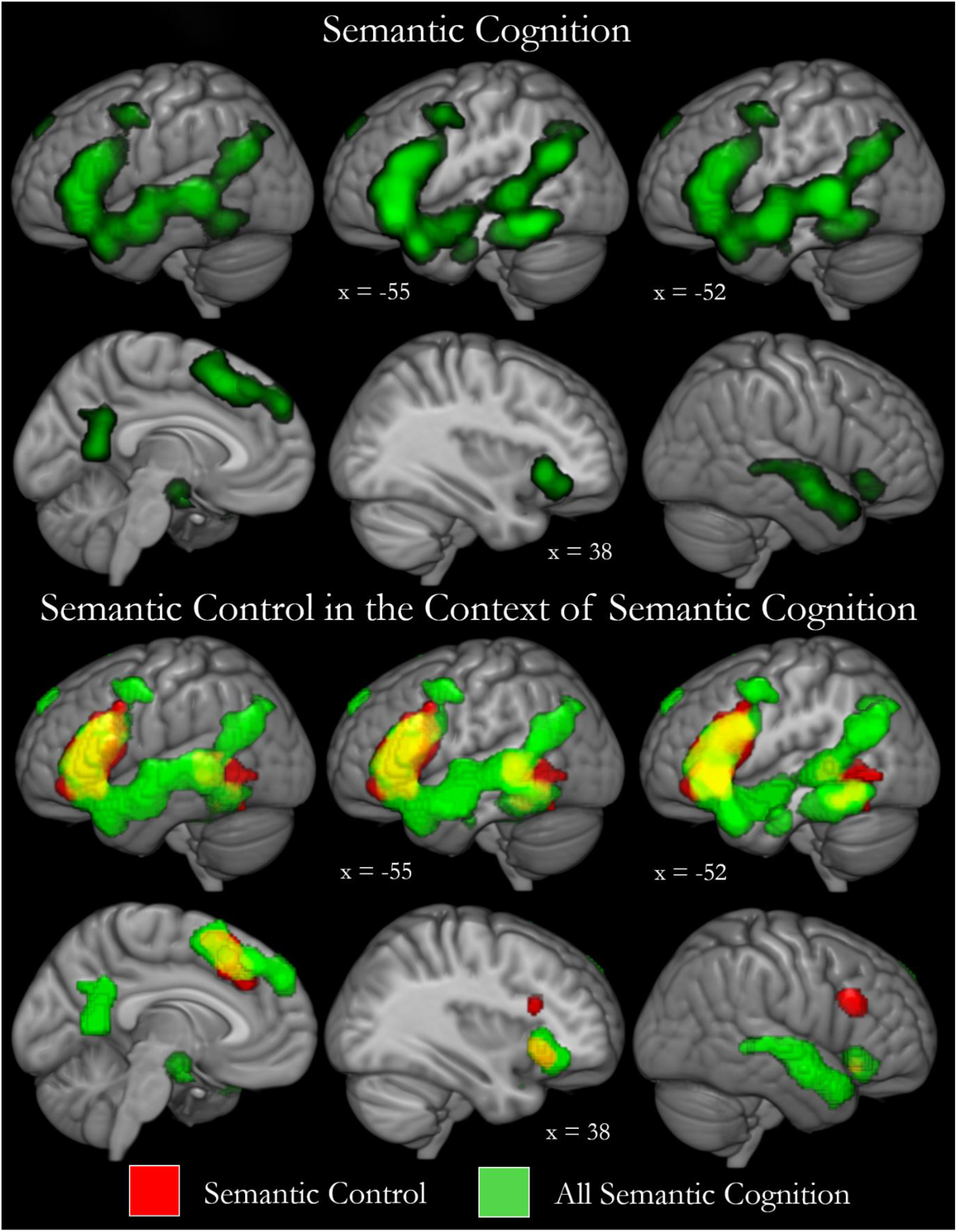
Semantic control in the wider context of general semantic cognition. Top: the regions reliably activated for semantic cognition are displayed. The semantic cognition meta-analysis contrasted semantic with non-semantic stimuli and tasks and includes 3860 peaks over 423 contrasts. Bottom: binary maps demonstrating how the semantic control regions fit within the wider network for semantic cognition.

**Table 3.**
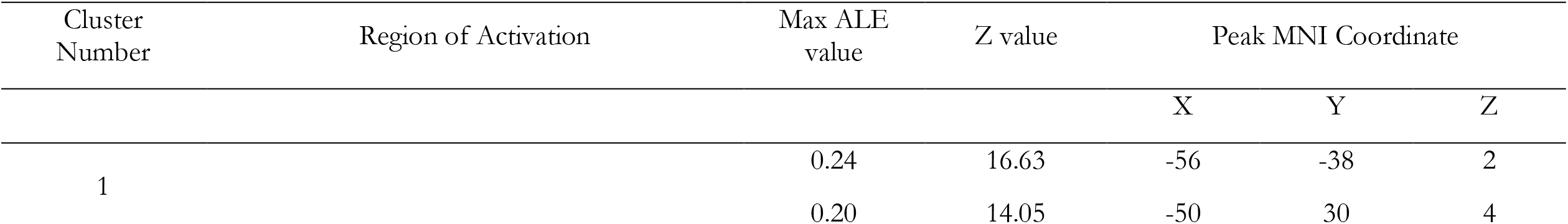
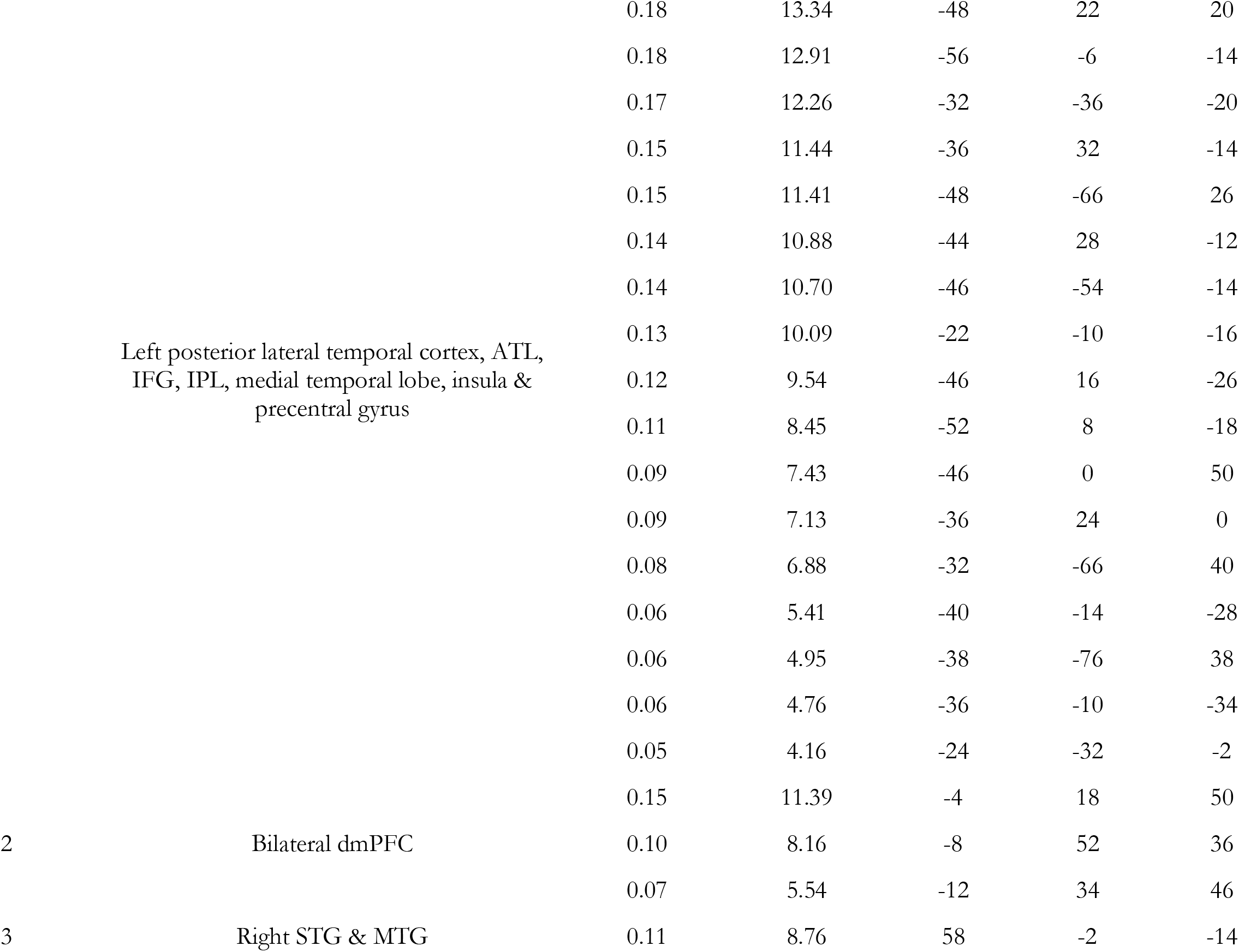
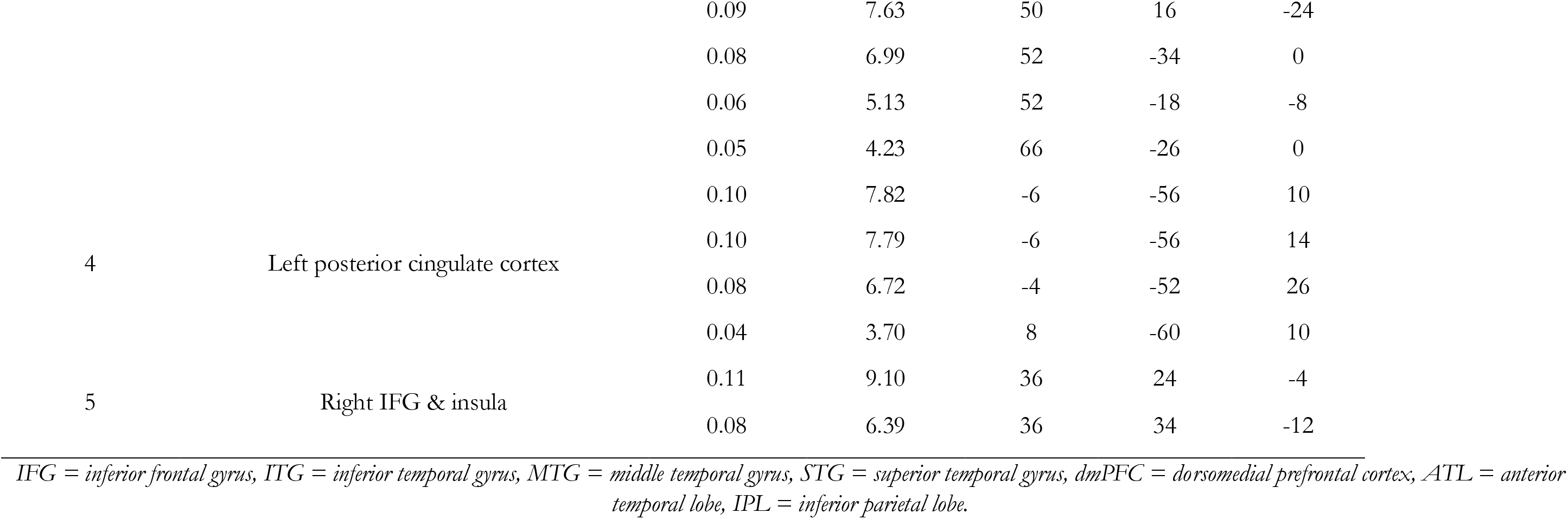
Semantic cognition activation likelihood.

## Discussion

Updated meta-analytic tools and a greater wealth of high-quality experimental data enabled a clearer picture of the topography of semantic control regions, in the context of the wider semantic network. Semantic control depends upon a distributed network consisting of IFG, posterior MTG, posterior ITG and dmPFC. This network is left-dominant with greater involvement of left than right IFG and no evidence for a role for right posterior temporal cortex. The networks found to underpin semantic control of auditory and visual stimuli were highly consistent, albeit with reduced involvement throughout for the auditory domain due to the lower number of eligible studies. Conjunction analyses were able to confirm the multimodal nature of the core network for semantic control, a key assumption of the Controlled Semantic Cognition framework which postulates that multimodal representation and control regions interact with modality-specific ‘spoke’ regions (Noonan et al., 2013; Lambon Ralph et al., 2017). The semantic control network operates in the context of a wider set of regions implicated in semantic cognition, found to additionally include the anterior temporal lobe and inferior parietal cortex. The meta-analysis results provide critical evidence to adjudicate upon three key puzzles within the anatomy of semantic control: 1) the involvement of, and topography across, inferior parietal cortex, 2) the extent of posterior lateral temporal involvement, and 3) the laterality of the semantic control network. Outside of these debates, the results are highly consistent with Noonan et al., (2013) with the improved statistical methods removing the small clusters in anterior temporal lobe (critical for semantic representation) and subgenual anterior cingulate (not implicated in semantic control, but may be recruited for particular aspects of semantic representation, such as emotional features; Etkin et al., 2011; Hiser and Koenigs, 2018). The rest of this Discussion addresses each of these puzzles in turn alongside key neuropsychological evidence and considers the potential next steps for semantic control research.

Unlike Noonan et al., (2013) the current, updated meta-analysis found no evidence for involvement of inferior parietal regions in semantic control. Here, there was greater statistical power and more appropriate statistical thresholding, therefore the previous IPL results could have been caused by a failure to account for multiple comparisons sufficiently. Alternatively, the difference may relate to the refinement in the inclusion criteria (e.g., the exclusion of studies that do not clearly contrast more over less controlled semantic cognition, such as comparison of metaphoric with literal meanings). The lack of ventral AG involvement in control is perhaps unsurprising. Identifying a region typically involved in easier than harder tasks for the opposing contrast was considered puzzling by Noonan et al., (2013). Although less surprising, the more dorsal AG/IPS cluster hypothesised to relate to domain-general control requirements may have reflected aspects of some studies tangential to the semantic control demands. Here, the AG is implicated in semantic cognition but not in control, suggesting a role in semantic representation. However, this ventral AG region was not the region most consistently identified in semantic cognition as in Binder et al.’s (2009) assessment. This may be due to a reduction in the difficulty difference between the semantic and baseline tasks (as the baseline tasks are no longer required to be at least as difficult) which would implicate default mode regions by virtue of their greater activation during less difficult task contexts (Humphreys et al., 2015), a possibility supported by the reduction throughout classical non-semantic regions of the default mode network (including the lack of significant findings in right AG and ventromedial prefrontal cortex). Alternatively, this may be due to the inclusion of nonverbal stimuli. The AG has been specifically associated with sentential and combinatorial processing (Graves et al., 2010; Humphreys and Lambon Ralph, 2014; Price et al., 2015; Solomon and Thompson-Schill, 2020; Branzi et al., submitted). Posterior SA patients typically have damage to large areas within temporal and parietal regions and therefore provide no clear evidence for a specific role for the parietal cortex.

As hypothesised, based on the spatial variability in peak activation within the literature, the involvement of lateral posterior temporal cortex in semantic control is more extensive than the pMTG alone. A large portion of both pMTG and pITG is implicated, bounded by the STS and with only a small region of fusiform gyrus reaching threshold. The term ‘pMTG’ may not be sufficient to describe the anatomy of the posterior temporal semantic control region and an alternative, such as ‘pMTG/ITG complex’ may provide a more transparent description of the particular anatomy of the region. Adequate localisation and labelling of this region is critical for understanding its role in semantic control, the interaction of control and representation regions, and the wider organisation of posterior lateral temporal cortex (associated with a large number of domains and semantic subdomains which could rely on the same underlying processes; see Kanwisher, 2017 for a review).

To date little research has explored how semantic representation and control processes interact; a complex issue due to their conflicting nature. Semantic representation requires the extraction of meaning that is preserved across contexts, whereas semantic control restricts behavioural output to be informed by context-relevant features only. A recent computational model demonstrated that the competing processes of semantic control and representation may co-exist within a system if its organisation promotes the relative specialisation of constituent regions for context-independent representations versus context-based responding (Jackson et al., submitted). In particular, the core demands of a semantic system were promoted only when the control signal interacted with shallower semantic representation regions (those closer to the modality-specific spokes than the multimodal hub). In neural terms this would equate to a prediction of no direct structural connection between the IFG control source and the ventral ATL hub (Jackson et al., submitted). Although this remains to be assessed, the low-level of long-range structural connectivity of the ventral ATL (Binney et al., 2012; Jung et al., 2016) strongly aligns with the possibility of an alternative, posterior route. One possibility is that connectivity between the IFG and the rest of the semantic system occurs via a multimodal control region in pMTG/ITG, well situated to interact with the visual and auditory regions in fusiform and superior temporal gyri respectively, before the representations become increasingly conceptual and multimodal in the progression anteriorly toward the ventral ATL hub (Binney et al., 2012; Davey et al., 2016). The role of pMTG/ITG as an intermediary between the frontal control and temporal representation regions would explain one further conundrum that has challenged the semantic control literature for the past decade; why does damage to inferior frontal and posterior temporal cortices result in the same behavioural profile?

Overall, the semantic control network was left-dominant, however the extent of this dominance varied by region. Whilst the dmPFC showed a bilateral pattern, clusters detected within right IFG were smaller and had a lower activation likelihood than within the left IFG. There was no evidence of right posterior temporal involvement in semantic control. Semantic cognition depends on a bilateral network, yet the regions recruited for a particular task vary based on multiple known factors, including the verbal or nonverbal status of the stimuli and the presence of visual or auditory stimuli, such that written words elicit the greatest left-dominance (Rice et al., 2015b; Rice et al., 2015a). Thus, the lateralisation within the semantic control network may also result from these factors, due to the almost exclusive use of verbal stimuli and the relative dominance of visual stimuli. This effect may be particularly strong in posterior temporal cortex if it engages in direct interaction with sensory-specific regions, which themselves vary strongly based on input type. Thus, it may be that manipulating the level of semantic control in nonverbal stimuli would shift the regions identified toward a more bilateral system and identify right pMTG/ITG. Alternatively, semantic control processes may truly be left-dominant within a bilateral semantic cognition system, making the necessary level of control an additional factor on which laterality of semantics-related activation varies. This possibility is supported by greater levels of intrinsic functional connectivity between left than right IFG and pMTG (Gonzalez Alam et al., 2019). Neuropsychological evidence may be able to distinguish these possibilities, however, the effect of right hemispheric stroke on semantic control is rarely studied. Thompson et al., (2016) identified a control impairment in a group of patients with cerebrovascular accident to right frontal or temporoparietal cortex that was qualitatively similar (but quantitatively reduced) than that of typical SA patients. However, the minority of participants had temporoparietal damage alone and the group analyses do not disentangle the specific roles of hemisphere and location.

Although not the focus of the current study, one further question is worth discussion; how does semantic control and its associated regions relate to domain-general control processes and topology? Several cortical areas have been postulated to perform control regardless of task domain, referred to as the multi-demand network (MDN; Duncan, 2010). Here, an inclusive definition of semantic control was employed, with the scope being to identify any regions responsible for control of semantic cognition regardless of their involvement across other domains. Thus, a high degree of overlap with the MDN is possible and perhaps even expected, yet relatively little is present, with the MDN centred primarily on more dorsal frontal and parietal cortices (Duncan, 2010; Assem et al., 2020). One clear exception to this is the dorsomedial prefrontal cortex (including supplementary motor and presupplementary motor area). This area is typically considered to have a general role, perhaps related to controlled motor output, consistent with its importance in speech production (Geranmayeh et al., 2014; Geranmayeh et al., 2017; Sliwinska et al., 2017). Although not a core region, the IFG is sometimes identified in multi-demand contrasts and recent assessments include a pITG region that may overlap the posterior temporal semantic control cluster (Duncan, 2010; Assem et al., 2020). Thus, the pattern appears to be one of relative differentiation with some shared substrates, suggesting further work directly contrasting these control processes is needed. Intriguingly, the majority of core MDN regions were not implicated in control of the wide range of tasks that employ meaningful stimuli, consistent with the observation that the frontoparietal control network may be disentangled from the networks recruited in semantic tasks (Jackson et al., 2019). The remarkable differences between the MDN and semantic control networks are consistent with comparisons between regions involved in language and domain-general control (Diachek et al., 2020), yet the focus on semantic control highlights a further subdivision. Whilst the regions implicated in domain-general control and semantic representation (including, but not limited, to verbal stimuli) do differ, a subset of regions relate specifically to the intersection of control and semantics (Davey et al., 2016). However, the differences between semantic and domain-general control regions (e.g., ventral and dorsal lateral frontal cortex) may be relative and a graded account may be best able to explain the pattern of cortical regions implicated in control and semantic cognition. Large convergence zones may perform control processes regardless of domain, yet the peak activation in these regions vary based on the location of structural connections to regions providing the subject matter for these computations (Assem et al., 2020). Such graded differentiation may underlie the posterior lateral temporal cortex, with semantic control demonstrating greater engagement of the pMTG and control of other domains preferentially engaging pITG. Further work is needed to disentangle the relations between domain-general and semantic control processes.

## Abbreviations

ALE: activation likelihood estimation
SD: semantic dementia
SA: semantic aphasia
IFG: inferior frontal gyrus
pMTG: posterior middle temporal gyrus
pITG: posterior inferior temporal gyrus
MTG: middle temporal gyrus
dmPFC: dorsomedial prefrontal cortex
ATL: anterior temporal lobe
MDN: multi-demand network
FWE: family-wise error
FDR: false discovery rate
AG: angular gyrus
IPS: inferior parietal sulcus
MNI: Montreal Neurological Institute
PET: positron emission tomography
fMRI: functional magnetic resonance imaging

## Acknowledgements

This work was supported by a British Academy Postdoctoral Fellowship awarded to RLJ (pf170068) and Medical Research Council intramural funding (MC_UU_00005/18). The funders had no role in the design, analysis or interpretation of the results or the writing of the manuscript. I am grateful to Dr. Rahel Schumacher for comments on the manuscript.

## Data & code availability

All data is included in the Supplementary Materials. Only freely available toolboxes were used to process the data. Masks of the results are available online.

## Ethics & competing interests

As no new data were collected, there were no ethical concerns. The author has no competing interests.

